# Evidence of an Unusual Poly(A) RNA Signature Detected by High-throughput Chemical Mapping

**DOI:** 10.1101/281147

**Authors:** Roger Wellington-Oguri, Eli Fisker, Mat Zada, Michelle Wiley, Eterna Players

## Abstract

Homopolymeric adenosine RNA plays numerous roles in both cells and non-cellular genetic material, and for lack of evidence to the contrary, it is generally accepted to form a random coil under physiological conditions. However, chemical mapping data generated by the Eterna Massive Open Laboratory indicates that a poly (A) sequence of length seven or more, at pH 8.0 and MgCl concentrations of 10 mM, develops unexpected protection to selective 2’-hydroxyl acylation read out by primer extension (SHAPE) and dimethyl sulfate (DMS) chemical probing. This protection first appears in poly(A) sequences of length 7 and grows to its maximum strength at length ~10. In a long poly(A) sequence, substitution of a single A by any other nucleotide disrupts the protection, but only for the 6 or so nucleotides on the 5’ side of the substitution. The authors are grateful for pre-publication comments; please use https://docs.google.com/document/d/14972Q36IDTYMglwMXTOrqd4P9orQ6-P3bPbCuITdv6A.

---

Homopolymeric poly(A) plays many regulatory roles in eukaryotes, prokaryotes^1^, organelles^2 3^, retroviruses^4^ and retrotransposons.^5^ Perhaps the most well-studied roles are those involving the lengthening and shortening of poly(A) tails added to various types of RNA sequences. In the case of mRNA, poly(A) tails are involved in transcription, nuclear export, translation initiation, protein synthesis regulation and, ultimately, decay of the mRNA.

Given all these roles, together with the observation that there is no information in a poly(A) sequence other than its length, it is reasonable to suspect that there is something about its structure that makes it so ubiquitous in vivo. Indeed, starting in the 1950’s, many experiments were done to determine the structure. Evidence was found of a single stranded structure forming under neutral pH,^6 7 8^ and specific molecular level single stranded models for it were proposed.^9 10^ More recently, new experimental techniques such as atomic force microscopy,^11^ nanoscale pores^12^ and vibrational circular dichroism^13^ have provided additional supporting evidence. Nonetheless, homopolymeric poly(A) still is generally assumed to form a random coil, or perhaps a stacked conformation like one strand of an A-form helix, at neutral pH.

Unlike previous experiments, chemical mapping provides measurements at the granularity of individual nucleotides. These measurements, in isolation, are not generally sufficient to determine RNA secondary or tertiary structure, but they can suggest molecular-level models testable by other experiments.

The data we report here come from the archives of Eterna.^14^ Eterna is an Internet-based citizen science project centered around designing, testing and analyzing synthetic RNA molecules. It is presented as a game, so participants refer to themselves as “players.” In Eterna’s first lab experiments, players created inverse folding puzzles — given a secondary structure, design an RNA molecule that adopts that folding. These secondary structures were often more challenging than naturally occurring RNAs.^15^ Players then voted on which puzzles would be tested experimentally, created submissions and voted for the best designs. A limited number of designs (typically ~100 per puzzle) were selected and given to the Stanford Das Lab, which performed chemical mapping experiments to determine the reactivity of each nucleotide of each design.^16^

Between 2013 and 2015, 27 rounds of high-throughput chemical mapping experiments tested more than 44,000 RNA sequences designed to solve over 500 unique puzzles. From these data, we have chosen puzzles that had single stranded sections of various sizes and loop type and also had a significant number of submissions with long poly(A) sequences. In order to eliminate any effect of variation in the experimental process over time, each of our examples is limited to data from a single experimental round.

Chemical mapping can be done with a variety of chemical probes, each measuring a somewhat different aspect of a nucleotide. SHAPE probes measure the reactivity of the 2’ hydroxyl group of the backbone, which is highly indicative of how flexible the backbone is at that nucleotide position and hence of base pairing.^17^ This makes the measurements comparable across all nucleotide types. DMS, on the other hand, reacts with N1 of adenosine and N3 of cytosine, while CMCT reacts primarily with N3 of U and N1 of G and detects the absence of nucleotide pairing.^18^ The SHAPE probe 1-methyl-7-nitroisatoic anhydride (1M7) was used in all of the Eterna experiments. In one round, three different probes with relevance to poly (A), 1M7, N-methylisatoic anhydride (NMIA), another SHAPE probe, and DMS were all used. In what follows, we report on the 1M7 data, adding the NMIA and DMS data where it is available.

Figure 1 shows the 1M7 measurements for a selection of designs that contain poly(A) sequences. This is the data display that first drew our attention to the unexpected protection of the 3’ end of a poly(A) sequence.

**Figure 1.**
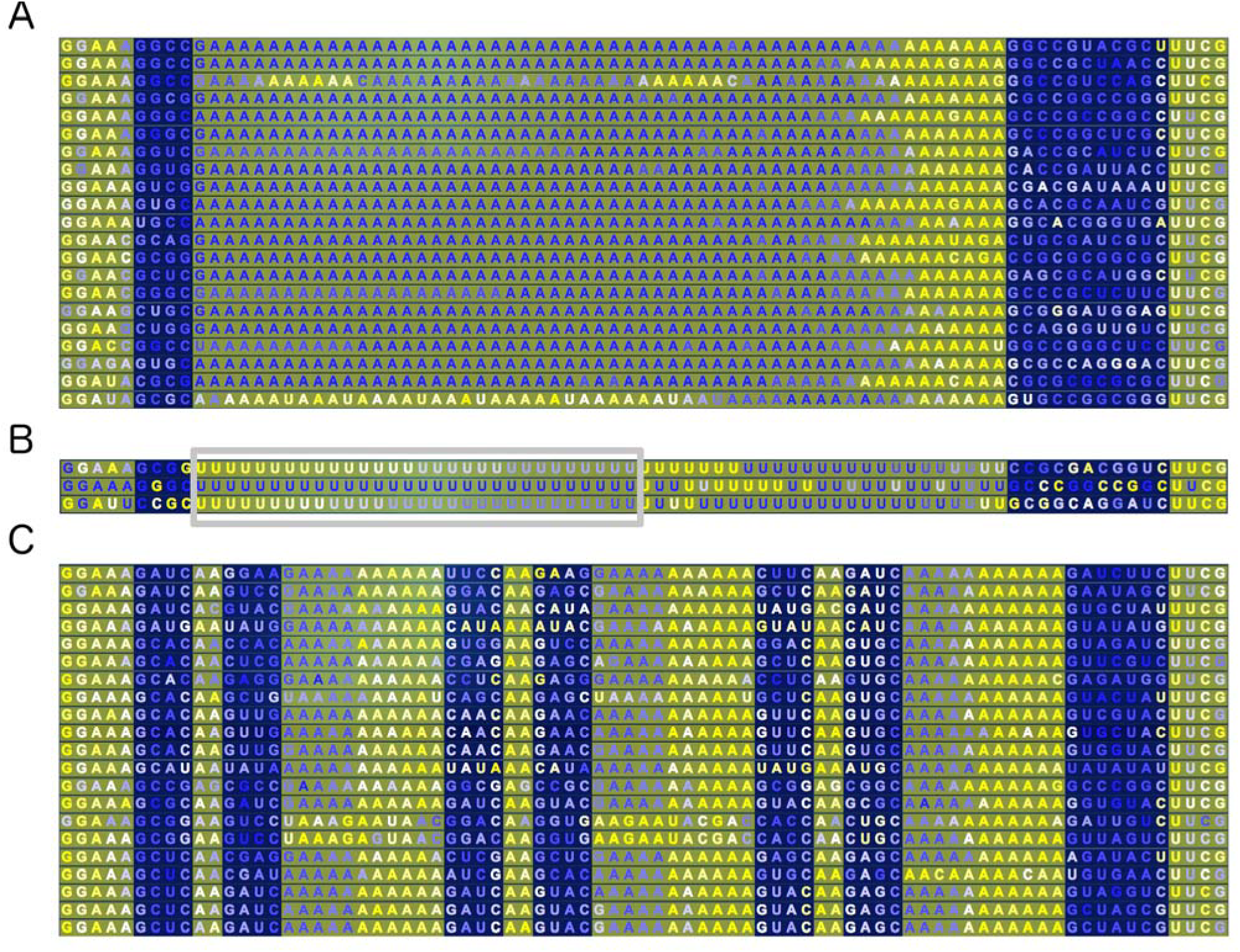
Excerpt of 1M7 SHAPE results for two lab puzzles. Nucleotide letter coloring varies from dark blue (highly protected) to bright yellow (highly exposed). Nucleotide background color reflects whether the puzzle expects the nucleotide to be paired (dark blue) or unpaired (grey.) (A) Big Hairpin Loop, designs selected for having 10+ consecutive unpaired adenosines. For most designs, only the ~6 adenosines at the 3’ end of the loop are fully exposed. Where a non-adenosine appears, the exposure is extended in the 5’ direction. (B) Big Hairpin Loop, designs with 10+ consecutive uracils. Despite having only three instances of poly(U), the pattern is clearly not at all like poly(A). The smooth gradient of protection values marked with a grey rectangle is an artifact of the limited sequencer read length used in the experiment. (C) Triangle of Doom, designs with 10+ consecutive A’s. The first two column ranges containing mostly A’s are hairpin loops. The third range is an exterior loop.

Figure 2 compares the median reactivity at each position of two subsets of the designs, for each of the puzzles in Figure 1. The first subset (left column) contains all those designs with only A’s over a defined range of positions. The second subset, for comparison, is all the designs that have no occurrence of more than six consecutive A’s.

**Figure 2.**
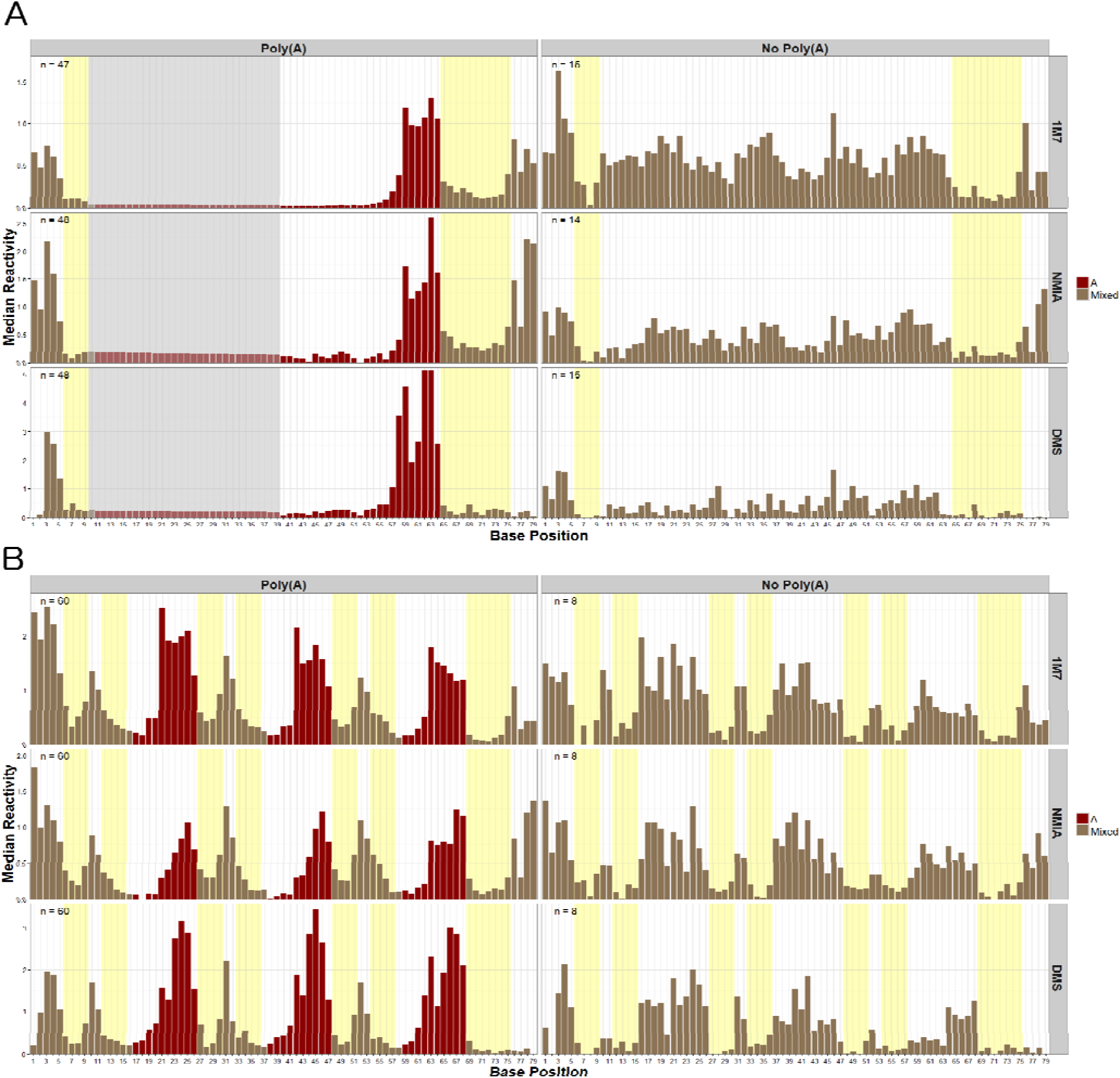
Chemical mapping values using three chemical probes, for the two puzzles in Figure 1. The left column displays the median value at each base position, over all designs that had adenosines at the positions colored red. The right column, for comparison, displays the median values over all designs for the same puzzle that had no more than 6 consecutive A’s, and shows a completely different distribution than seen in the left column. The number of designs satisfying the respective criterion is shown in the upper left corner of each graph. Experimental conditions were pH 8.0 and 10nM MgCl for all three probes. (A) Big Hairpin Loop. The specified A sequence is 44 nucleotides of a hairpin loop of length 45. The grey background indicates positions where reactivity cannot be assigned at single nucleotide resolution, due to experimental limitations. (B) Triangle of Doom. The specified A sequences are of length 12 in each of three separate loops, two hairpins and one exterior.

The experimental round for this data tested all the designs with three relevant chemical reagents 1M7, NMIA, and DMS. NMIA, like 1M7, is a SHAPE probe that forms a chemical bond with the 2’ hydroxyl of the back-bone sugar. In contrast, DMS methylates the N1 of adenosines (and N3 of cytosines).

In Figure 2, the two SHAPE probes, 1M7 and NMIA, show a similar pattern of decreased exposure for all of the poly(A) nucleotides except the six 3’-most ones. But the distribution of exposure values at the 3’ end of the loops to 1M7 and NMIA is also noteworthy, especial in figure 1B. Differences in reactivity between these two SHAPE reagents have been proposed as a way to detect backbone configurations that are different than those found in A-form double helices.^19^

Overall, the DMS exposure closely follows the 5’ versus 3’ distinction of the two SHAPE probes. To the best of our knowledge, there is no report in the literature of adenosine being protected from DMS other than by having N1 forming a hydrogen bond. This implies that the 5’ A’s may be forming hydrogen bonds

But in contrast to the SHAPE probes, high DMS exposure occurs for seven, rather than six, of the 3’-most nucleotides in the Big Hairpin Loop and in the open loop in the Triangle of Doom. This difference in protection for the seventh nucleotide suggests that the protective constraints on the backbone and the adenine components of the putative structure do not rely on the identical mechanism. This is consistent with a structure that relies on the combination of Mg^+2^ stabilization of the backbone and hydrogen bonding of the adenines.

Finally, although the general 3’ vs 5’ distinction seems to be universal, Figure 1B does indicate that the secondary structure surrounding the poly(A) sequence does have some effect. In the left column, one can see differences between the two poly(A) hairpins (which are very similar to each other) and the exterior loop poly(A) strand. This is particularly noticeable in the NMIA data.

So far, we’ve only looked at poly(A) lengths that are long enough for the protection pattern to be present. In order to determine the minimum length of poly(A) for the pattern to emerge, we turned to an Eterna round that tested 104 different target shapes, for the purpose of stress-testing the high-throughput chemical mapping method developed to replace capillary electrophoresis^16 20^. This round was chosen because the large number of puzzles created a wide range of single stranded sequence lengths. Figure 3 summarizes the pattern of protection from 1M7 for poly(A) sequences of lengths from 5 to 14.

**Figure 3.**
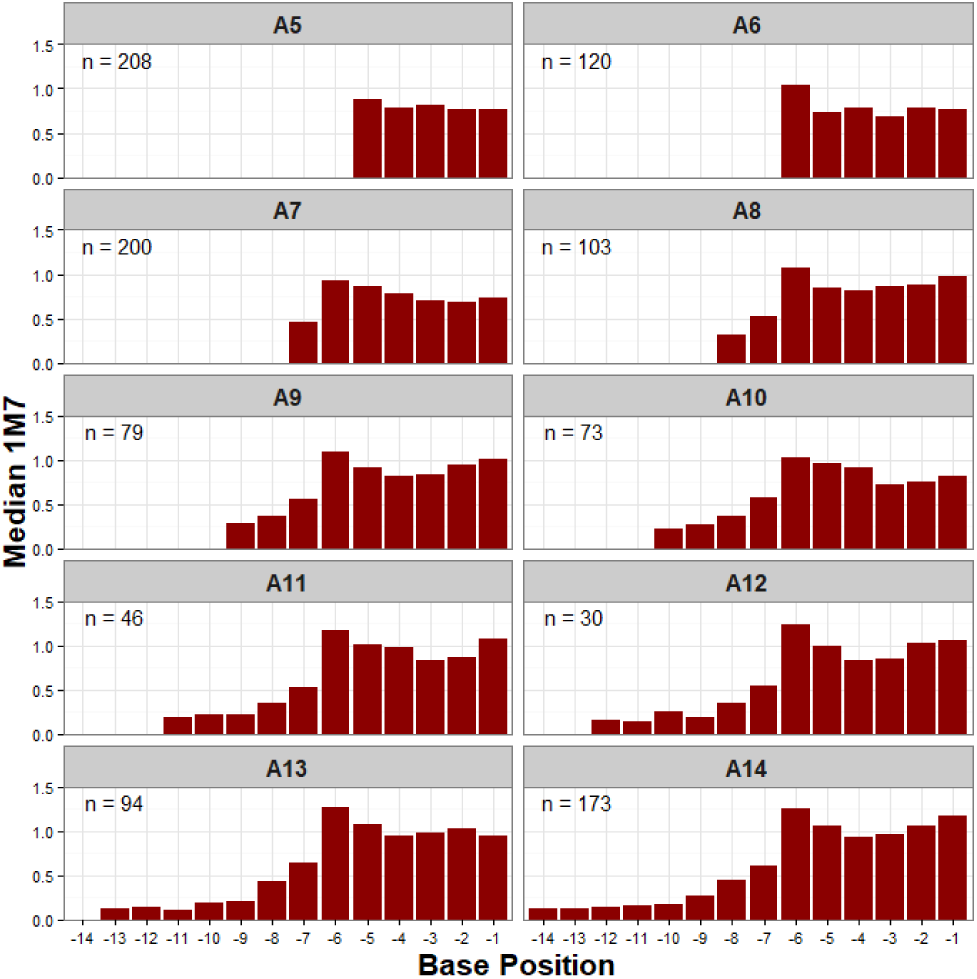
Protection from 1M7 of poly(A) sequences of length 5 to 14 nucleotides. Base positions are numbered relative to the non-A following the 3’ end.

The reactivity of the individual nucleotides is near constant for poly(A) sequences of length five and six. However, starting with length seven sequences, all but the six 3’-most nucleotides show some protection from the 1M7 probe. As the length increases, the A at the added position is even more protected, with the decrease approximating an exponential curve.

The final question we considered is what happens when a long poly (A) stretch is interrupted by a single non-A base? Does the choice of interrupting base make a difference? To address this, we examined the data from a puzzle that was specifically proposed by one of the authors to answer that question. In this puzzle eight out of each sequential nine nucleotides were required to be A, with the ninth being chosen by the sequence designer. The signal-to-noise ratio for this synthesis round was lower than usual but graphing the median values of the large number of sequences submitted allows a pattern to stand out. The results are shown in Figure 4.

**Figure 4.**
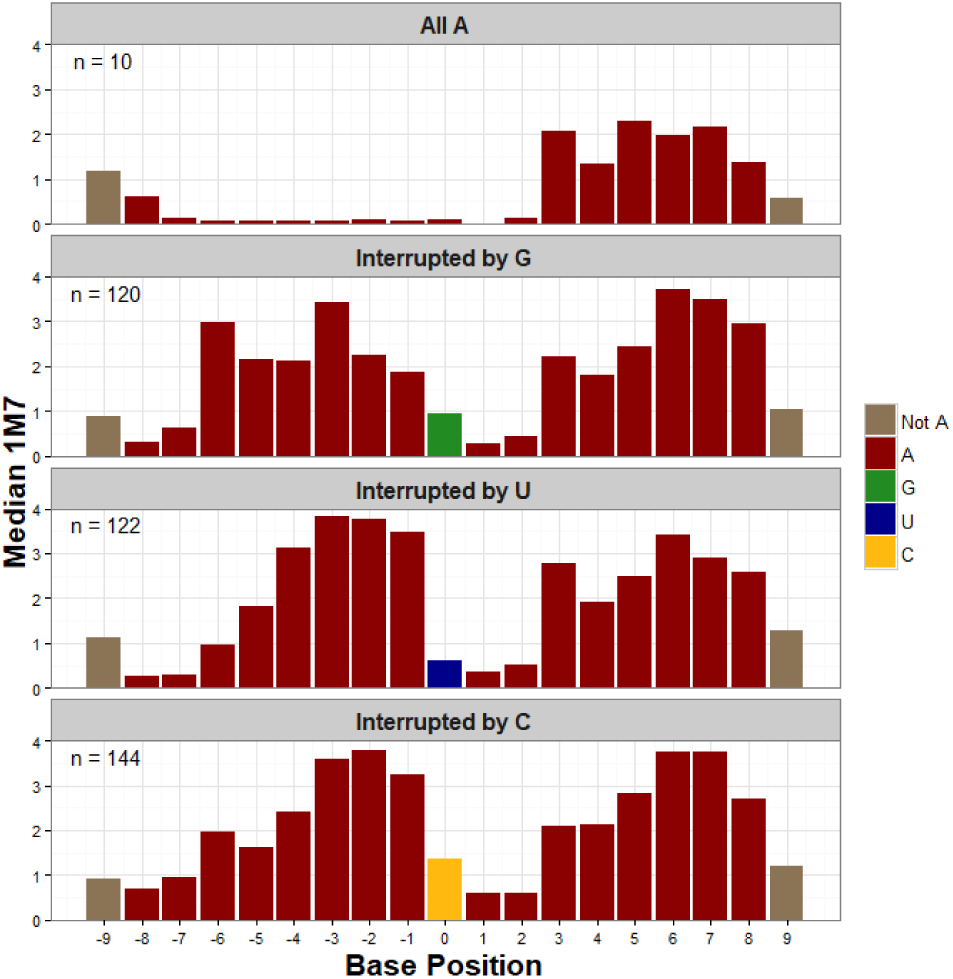
Results from Intrinsical - Frequency 8 puzzle. Protection (1M7) of A17, interrupted by a single non-A nucleotide in the middle position. Base positions are relative to the middle position.

The top graph shows the degenerate case, where the center nucleotide is another A. There were only 10 instances, but the result is consistent with the protection pattern described previously. Each of the other choices for the center nucleotide clearly separate the sequence in two, with each 8-nucleotide sequence showing the expected 6 3’-most nucleotides more exposed and the two 5’ nucleotides more protected.

The bottom three graphs also show some interesting differences among themselves. The specific choice of interrupting nucleotide does not appear to have a large effect on the (two) nucleotides at the 3’ end. However, they clearly have a differential effect in the 5’ direction. Here, the reactivity falls off faster for the two pyrimidine nucleotides, C and U, than for the purine G. If we envision the interfering nucleotide as a defect introduced into the structure, it is as though the smaller purine nucleotides create a smaller defect, allowing the structure to reform slightly more quickly.

In summary, the data suggest that poly(A) sequences of length seven or more do form a well-defined structure at pH 8.0 and MgCl concentrations of 10 mM. The structure requires a minimum of seven consecutive A’s, and in longer sequences, a clear difference arises between the 5’ and 3’ ends. The 5’ end becomes very resistant to chemical interaction, while the six adenosines at the 3’ end become more reactive than those in a single stranded sequence of six or fewer A’s. This structure seems to be dependent on both hydrogen bonding between the nucleic acids and stabilization of the backbone by Mg^+2^ ions; however, we cannot rule out configurations whose chemical reactivity to DMS or SHAPE reactions is perturbed by other effects, such as electronic rearrangements that may be associated with syn conformations of the nucleobase. Finally, a single non-A nucleotide is sufficient to interrupt the structure, but it forms independently on each side of the interruption.

Because the putative structure for poly(A) would have such a large effect on subsequent biological research, the Das lab has been conducting additional gel electrophoresis experiments to verify that the data are not artifacts of the massively parallel experimental process.^21^ These experiments have confirmed the details presented here. However, experiments haven’t yet ruled out one last possibility -- that the reverse transcriptase has a previously undiscovered ability to consistently transcribe even chemically modified A’s once it has transcribed ~6 consecutive unmodified A’s. A new experiment is currently being conducted that should rule out or confirm that hypothesis.

Why is poly(A) so ubiquitous in nature? An understanding of this structure at atomic resolution may lead to advances in understanding what are the unique characteristics of poly(A).

As one example, consider the addition of the poly(A) tail to most eukaryotic mRNAs. It is known that after the cleavage and polyadenylation specificity factor (CPSF) binds to a pre-mRNA, a polyadenylate polymerase (PAP) can start creating the poly(A) tail. However, the tail grows distributive because the PAP is loosely bound and falls off the growing tail after each addition. Once the tail has grown to a length of 10 or 11 nucleotides, nuclear poly(A) binding protein (PABPN1) binds to the poly(A) complex, and the tail grows rapidly to its mature length.^22^

X-ray crystallography has determined the atomic level structure of the PABPN1 RNA recognition motif (RRM) that serves to bind the PABPN1 selectively to the sufficiently long poly(A) tail.^22^ However, the atomic level details of how the RRM binds to the poly(A) is still a mystery.^23^ Determining the structure implied by our data is likely to answer this question.

Another open question, involving templated poly(A) sequences, is why mRNA translation stalls when polylysine is encoded for by three consecutive AAA codons, i.e. nine consecutive A’s.^24^ Since this put the number of consecutive A’s in the range where the inferred structure forms, it may well be part of the causal chain.

These are but two examples where it seems that the character of poly(A) changes as the length transitions through the 7–12 range. Given the ubiquity of poly(A) in nature, there are likely to be many others. We hope calling attention to the chemical mapping data will help spark interest in understanding what is happening and why poly(A) seems to be involved in so many different biological activities.

## Author Contributions

The manuscript was written through contributions of all authors. All authors have given approval to the final version of the manuscript.

## ACKNOWLEDGMENT

We thank Rhiju Das for encouragement to organize our observations into a form suitable for publication and for his generous editorial guidance.

## ABBREVIATIONS

SHAPE, Selective 2’ hydroxyl acylation analyzed by primer extension; NMIA, N-methylisatoic anhydride; 1M7, 1-methyl-7-nitroisatoic anhydride; DMS, Dimethyl sulfide, PABPN1, Nuclear poly(A) binding protein 1; PAP, poly(A) polymerase; RRM, RNA recognition motif.

